# Linking scales of life-history variation with population structure in Atlantic cod

**DOI:** 10.1101/2020.07.17.208660

**Authors:** P.J. Wright, A. Doyle, J.B. Taggart, A Davie

## Abstract

It is increasing recognised that sustainable exploitation of marine fish requires the consideration of population diversity and associated productivity. This study used a combination of genotypic screening and phenotypic traits to define the scale of population structuring in Atlantic cod inhabiting the northern North Sea (ICES 4a) and Scottish west coast (6a). The genetic analysis indicated an isolation by distance pattern with an even finer scale structuring than previously reported, that persisted over a decade and between feeding and spawning seasons. Spatial variation in phenotypic traits reflected genetic variation with cod maturing later and at a larger size near the Viking Bank in 4a. The identified population structuring provides an explanation for differences in historic changes in maturation schedules and the temperature exposure recorded in previous electronic tagging studies. The study also highlights how the mismatch between stock divisions and population units is leading to a misunderstanding about stock recovery.

## 1. INTRODUCTION

Conserving population diversity is a key part of the UN Convention on Biological Diversity as it promotes persistence and adaptability in taxa (Balmford et al., 2005). Although the need to account for intraspecific genetic diversity has been widely acknowledged (Hilborn et al., 2003; Ruzzante et al., 2006) and overfishing has been related to a decline in genetic diversity across a wide range of marine fishes (Pinsky & Palumbi, 2014), it is still rarely considered in marine fisheries management. This is because the stock units upon which management advice is based often do not match the scale of population structuring (Stephenson, 2002; Barth et al., 2017). While these stock units are assumed to be a geographically discrete group of fish with the same vital rates, they usually reflect a compromise between biological evidence, data availability and management requirements (Stephenson, 2002). Failure to recognise the appropriate scale of population processes may mask changes in the sub-stock components and lead to biased estimates of abundance and overexploitation of less productive populations (Smedbol & Stephenson, 2001). Accounting for within-stock differences in productivity is far more important to management than just the knowledge of population structuring, because of the assumption of homogeneity in vital rates (Stephenson, 2002). Despite advances in the genetic and tagging tools available, few studies have related genetic differentiation with vital rates in marine fish. Consequently, in order to provide advice on conserving population diversity and sustainable exploitation in marine fish, there is a need for studies that characterise both population distribution and productivity.

Reproductive traits have often been used as a proxy of populations’ vital rates or productivity as they are shaped by regional variation in growth and reproductive success while also responding rapidly to local differences in environmental conditions (Pianka, 1976; Wright, 2014). Accurate information on maturity at age is also needed for analytical assessments to estimate spawning stock biomass and biomass reference levels. While inter-annual variation in maturity-size relationships are affected by growth rate (Godo & Moksness, 1987) and temperature (Tobin & Wright, 2011) there have also been substantial downward shifts in maturation reaction norms that are related to historical fishing pressure rather than either of these natural factors (Devine et al., 2012). Consequently, a move to conserving population diversity requires information on the appropriate scale of variation in maturity at size and age.

Atlantic cod, *Gadus morhua*, is well suited to studying population diversity and reproductive traits as there is already considerable information on both genetic divergence (Bradbury et al., 2013; Barth et al., 2017) and variation in maturity and fecundity (Wright & Rowe, 2019). Like most marine fish the level of genetic differentiation based on unlinked neutral SNPs appears to be very low in Atlantic cod (Bradbury et al. 2013; Berg et al., 2015) with most reported genomic divergence being related to polymorphic chromosomal rearrangements linked to adaptive loci (Bradbury et al. 2013; Berg et al. 2015, Sodeland et al., 2016). However, these genomic regions of adaptive divergence appear very relevant for advising on the appropriate scale for stock management as they appear to reflect reproductive barriers linked to local environmental adaptation and migratory phenotypes (Bradbury et al. 2013; Barth et al., 2017; Kess et al., 2019). Variations in reproductive traits at scales less than the fished stock have been widely reported in this species (Marteinsdottir & Begg, 2002; Yoneda & Wright, 2004), although only in Norwegian coastal cod has this variability been related to genetic differences (Olsen et al., 2009). Downward trends in reaction norms towards smaller and earlier age at maturation have been widely reported for many cod stocks (Devine et al., 2012), and regional differences in these trends within the North Sea have been linked to historic differences in fishing effort (Wright et al., 2011).

In the North Sea (International Council for the Exploration of the Sea (ICES) 4) and off the Scottish west coast (ICES 6a), studies of single nucleotide polymorphic (SNP) markers of spawning cod provide evidence of barriers to gene flow between the deeper north east North Sea region between 100 – 200 m and the shallower shelf region throughout these two stock areas, except for the Clyde Sea (Heath et al., 2014). Analyses of otolith chemistry, comparing larval, juvenile and adult parts for the same year-class in the North Sea, suggests that this reproductive isolation is maintained by a combination of hydrographical isolation of early life-stages and either fidelity or natal homing of later stages (Wright et al., 2006a; Svedäng et al., 2010; Neat et al., 2014; Wright et al., 2018). Spatio-temporal differences in maturity at size seem to be consistent with the scale of mixing suggested by tagging (Wright et al., 2006a; Neat et al., 2014), with differences found among cod in the north west, north east and southern North Sea and Scottish west coast (Yoneda & Wright, 2004; Wright et al., 2011). However, small differences in the geographic region selected can have a substantial impact on maturity at size, as evident from differences reported for the north east (Wright et al., 2011) and northern North Sea (Heath et al., 2014). Hence, there is a need to examine how reproductive traits co-vary with genetic variation.

In this study, we examine how variation in growth and maturity relates to population scale in Atlantic cod across two stock areas; the North Sea (ICES 4) and Scottish west coast (ICES 6a). Building on Heath et al. (2014) and the emerging focus on chromosomal rearrangements, we extend the range of sampled locations to better define population boundaries across the region and consider genomic variation at both spawning and feeding times in 2013-14 and compare with samples collected in 2002-3 including some samples used in Heath et al. (2014). Regions of genetic discontinuity are then compared to the spatial variation in length and maturity at age. By this means we test whether variation in reproductive traits are linked to population diversity.

## 2. MATERIALS AND METHODS

### 2.1 Sampling

As cod are known to aggregate to spawn (Wright & Rowe, 2019) but tend to disperse when feeding, we sampled in autumn, shortly after the time they allocate energy to secondary gametogenesis (November – December), and thereafter during spawning (February – March). A total of 721 cod were sampled in November to December in 2013, when early stages of oogenesis could be detected histologically (Yoneda & Wright, 2005) and a further 803 during the spawning season between February and March (Gonzalez-Irusta & Wright, 2016) in 2013 and 2014. Of these 1524 fish, a sub-sample of 658 were selected from six locations and the two periods for SNP identification (Figure 1). Cod were sampled by bottom trawl from the region 57°N to 61°N and 7°W to 5°E from both commercial and research vessels. A further sample of 386 cod collected over the same region in February – March 2002-3 (n=254) and August – November 2003 (n= 132) for an earlier study (see Heath et al. 2014) were used to provide a temporal comparison for genetic analysis.

**Figure 1.**
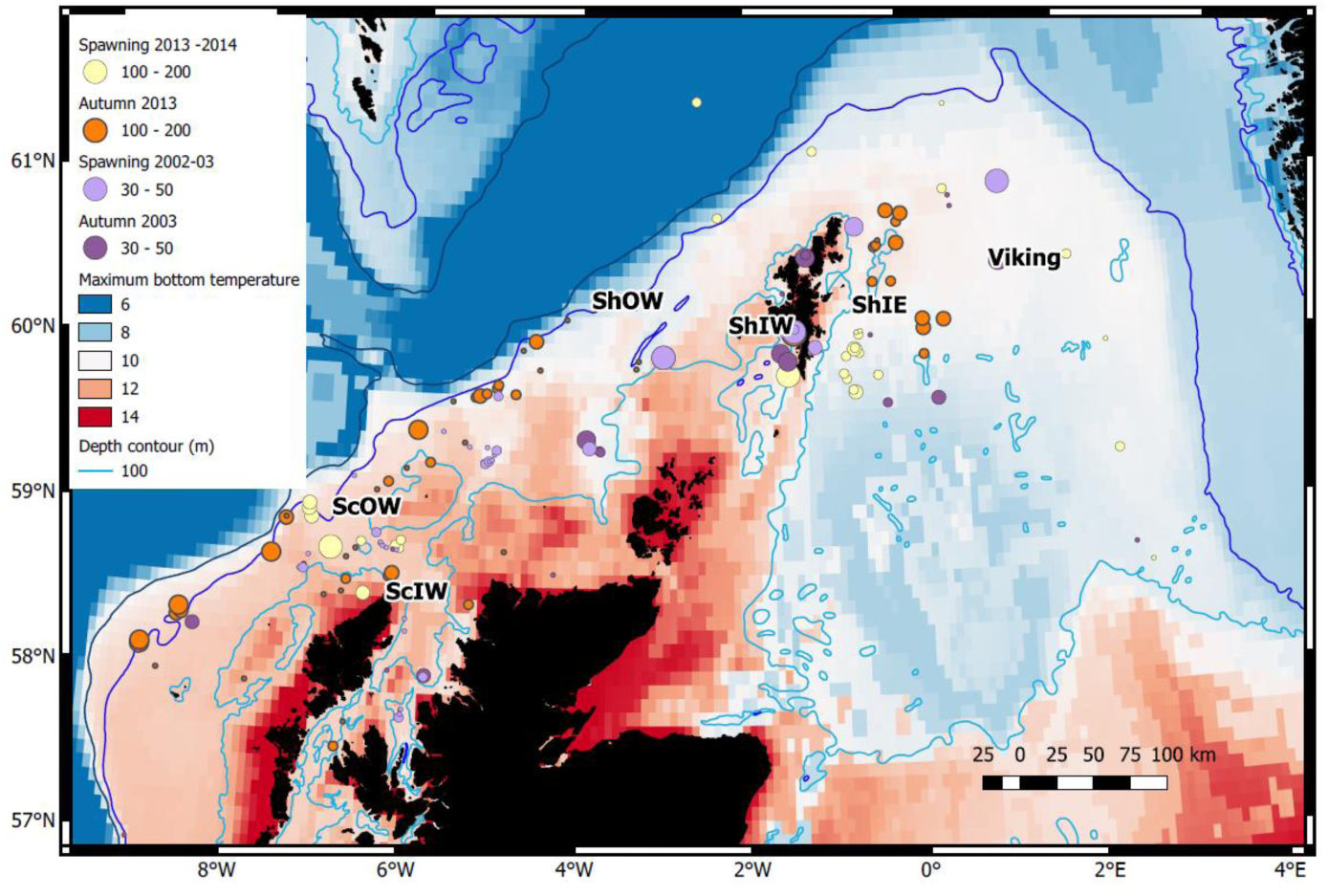
Distribution of genetic samples in relation to maximum seasonal bottom temperature (°C) for the spawning and autumn seasons in sampled years. The centre of 6 areas identified in Table 1 are indicated. Circle diameters are related to sample size (note larger sample sizes in 2013-14). The 200 m depth contour is also shown.

All cod > 11 cm TL were sampled for sex, maturity stage and total length (± 0.5 cm). Sagitta otoliths were taken for all samples for annual age estimation. For all cod from the February/March data set and all male cod from the November-December dataset, maturity stage was determined macroscopically according to ICES guidelines (Bucholtz et al., 2007). Female cod (*N* = 418) from the November/December samples were staged histologically to ensure that early maturation commitment, evident from oocytes in cortical aveoli and vitellogenesis stages (Yoneda & Wright, 2005), was identified. Ovary tissue was fixed and stored in 10 % neutral buffered formalin solution before being embedded in paraffin wax, sectioned at 2μm using a rotary microtome RM2155 (Leica Instruments GmbH) and stained with haematoxylin and eosin. Gill clips from sampled fish were preserved in 100% ethanol for the SNP analysis.

**Table 1:**
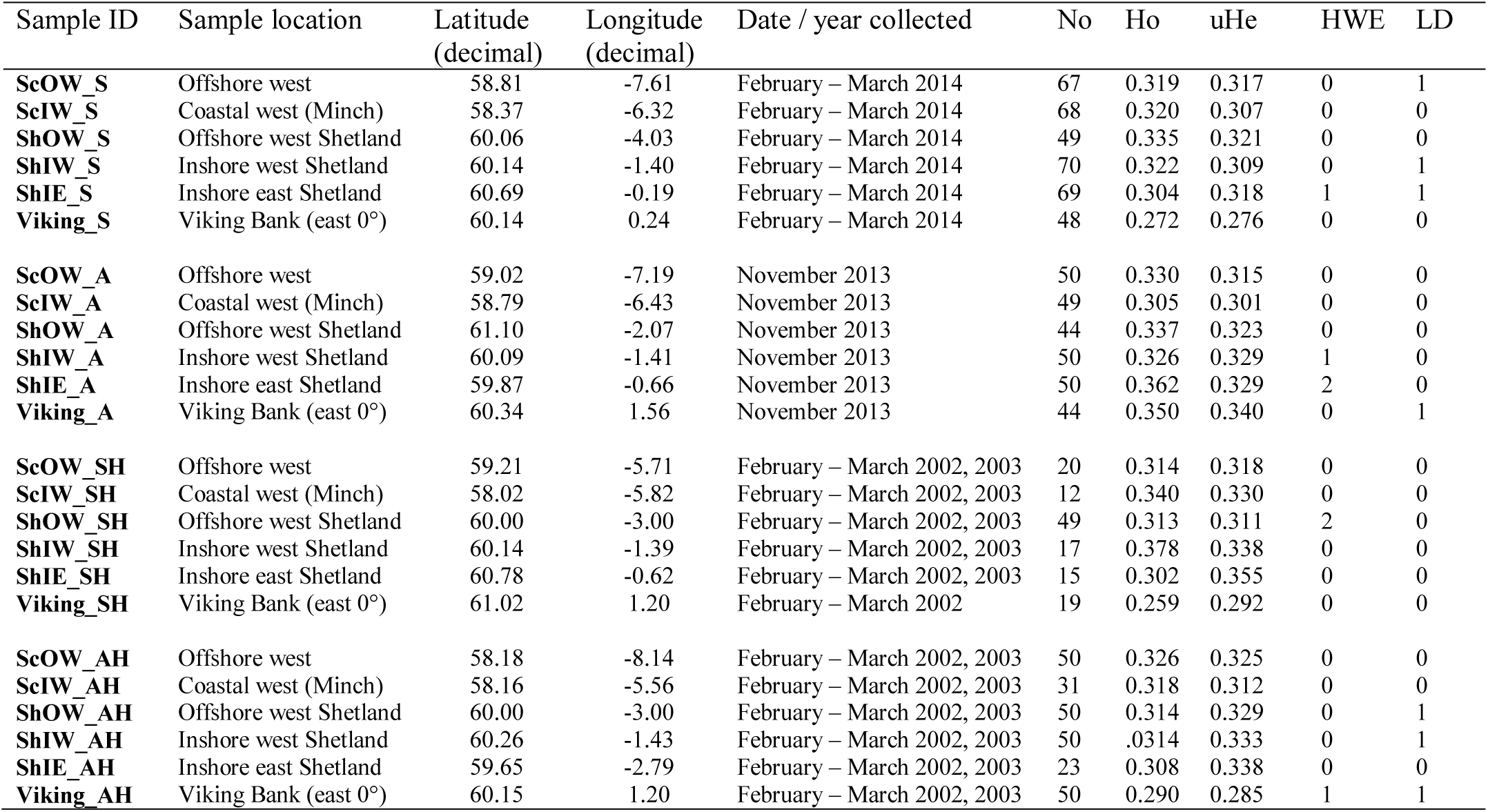
Overview of sample origin, and summary of population specific SNP marker (n = 13) performance listing unbias observed (Ho) and expected (uHe) heterozygosity, as well as the number of markers which deviate from Hardy-Weinberg equilibrium (HWE) or Linkage Disequilibrium (LD).

In order to model maturity at size variation, the two spawning datasets were supplemented with similar data extracted from the 1st quarter ICES International Bottom Trawl surveys (http://ices.dk/marine-data/data-portals/Pages/DATRAS.aspx); the North Sea International Bottom Trawl Survey (NS-IBTS; 2002-3 and 2013-2014), the Scottish West Coast Bottom Trawl Survey (SWC-IBTS; 2002-3) and the Scottish West Coast Groundfish Survey (SCOWCGFS; 2013-2014). The NS-IBTS takes place in January – February whilst the Scottish West Coast surveys are in February – March. Combined with the dedicated sampling this provided a sample size of 10,139 cod.

### 2.2 Growth and Maturity analysis

Variation in total length (L, mm) for a given age in relation to decimal latitude and longitude was examined with the addition of sex and year, treated as factors. A generalized additive model (GAM) with a Gaussian distribution was used to account for the non-linear trends with latitude and longitude. Model selection was made by forward selection of variables based on a significant decrease in residual deviance and reduction in the Akaike information criteria (AIC).

Spatial variation in maturity (M) was modelled using a binomial GAM with a logit link according to:

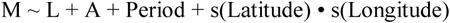

where M is the proportion mature (0/1), and the two periods of study (2002-3 and 2013 – 14) were treated as a factor. Sexes were examined separately given previous evidence for sex related differences in maturity at size (Wright et al., 2011). The same model was examined for between season differences in female maturity, but with the period term representing the feeding and spawning sampling times in 2013 – 14. Due to over dispersion in the survey data, a quasi-binomial link function was used where variance is given by the dispersion parameter multiplied by the mean. As there were few cod aged >5, all analyses were restricted to ages 2 – 5. Model selection was made by forward selection of variables based on a significant decrease in residual deviance. All models were implemented in R3.3 using mgcv and MASS libraries.

### 2.3 DNA Extraction and RAD Library Construction

DNA was extracted from ethanol preserved gill samples using a salt extraction method and treated with RNase to remove residual RNA from the sample. Each sample was quantified by fluorimetry (Quibit) quantitation and quality assessed by agarose gel electrophoresis, and was finally diluted to a concentration of 50 ng.μL^-1^ in 5 mmol.L^-1^ Tris, pH 8.5.

### 2.4 SNP identification

In order to identify a reduced panel of informative SNPs, adopting a similar approach to that used in recent studies of a nearby geographic region (Barth et al., 2017), DNA samples from 20 individuals from each of the six locations from the contemporary February – March sampling window were taken forward into RAD analysis. A double digest RAD library (Peterson et al., 2012) was constructed according to the methodology of Brown et al. (2016). Briefly, the restriction enzymes Sbf1 and Sph1 were used to digest 24 ng of genomic DNA from each sample, followed by ligation of individual-specific combinations of P1 and P2 adapters, allowing subsequent post-hoc segregation of all samples. The library was run twice on an Illumina MiSeq platform (v2 chemistry, 150 base paired-end reads). Data were compiled and processed using STACKS v1.27 (Catchen et al., 2013). Following data processing a candidate list of robust, biallelic loci were taken forward into initial marker description, including Hardy-Weinberg equilibrium and pairwise population F_st_ computations performed in GENEPOP on the Web v 4.2 (Raymond & Rousset, 1995; Rousset, 2008), while ARLEQUIN v 3.5 (Excoffier & Lischer 2010) was used to examine for signatures of directional selection by implementing a hierarchal island model (20,000 simulations, 100 demes per group, 10 groups). Following this screening, 90 ‘neutral’ loci with a minor allele frequency ≥ 0.15 in at least one of the test populations and an Fst ≥ 0.03 were identified as being putative loci which could be informative for population structuring in the study range. The candidate list was further rationalised to 13 loci (Supplementary Table S1) based on the potential resolving power in relation to the study areas (F_st_ value), confirmation (blastn) of sequence alignment to a single high identity return within the cod genome (gadMor1), screening and removing loci with tandem repeats evident within the 150 read (Benson, 1999) and ensuring that a minimum of 50 bases flanking sequence either side of the SNP was present to ensure that the sequence meets the requirements for the design of a genotyping probe assay (LGC Genomics, 2014). These SNPs were translated into genotyping assays designed using the KASP on demand genotyping system (KASP™ v4.0, LGC Genomics, UK), and a total of 1044 samples (Table 1) were genotyped by LGC genomics.

### 2.5 Genetic analysis

For the full data set, allele frequencies and unbiased expected (uHe) and observed (Ho) heterozygosity were estimated using the package GenAlEx 6.5 (Peakall & Smouse 2006, 2012). Departure from Hardy Weinberg equilibrium (HWE) and linkage disequilibrium (LD) for each pair of SNPs was tested in each population using GENEPOP on the web (Raymond & Rousset, 1995; Rousset, 2008). Type I error rates were corrected for by using a sequential Bonferroni procedure for both tests. Genetic differentiation was tested using the Analysis of Molecular Variance (AMOVA) using GenAlEx 6.5 (Peakall & Smouse 2006; 2012) as well as by pairwise Fst using GENEPOP on the web. Bayesian clustering analysis (STRUCTURE v2.3.4) was performed using an admixture model and correlated allele frequencies among populations, as well as providing sampling information as a prior in order to improve accuracy in detecting population structure. Results were compiled using CLUMPAK (Kopelman et al., 2015). The dataset was also processed using a discriminant analysis of principal components using the ADEGENT program in R (Jombart, 2008, Jombart & Ahmed, 2011). Geographic distance among sampled areas shown in Figure 1 were calculated in QGIS using the DDJOIN plugin.

## 3. RESULTS

### 3.1 Spatial variation in length at age

Location, in terms of latitude and longitude, explained the highest model deviance in length for all age-groups (Table 2), with a decline from south west to north east (Figure 2). There was no linear trend in length variation, except for the effect of latitude on age 3 cod. A difference in length among sampled years was only found for age 2 cod (p = 0.02). Sex had a significant effect on length for ages 3 and 4 across the study area (p < 0.01), with males being slightly shorter at age. No interactions were significant and none of the models explained a high percentage of the deviance, with the highest being for age 3 cod.

**Table 2.** GAM model coefficients, deviance explained and AIC for length in relation to year, longitude, latitude and sex for ages 2-4.

|  | Coefficients |  |  |  |  |
| --- | --- | --- | --- | --- | --- |
| Explanatory | Estimate | SE | *p* | Deviance (%) | AIC |
| **(a) Age 2** |  |  |  |  |  |
| Intercept | 379.396 | 3.303 | <0.001 |  |  |
| Year | 10.427 | 4.506 | 0.0209 | 2.06 | 8816.335 |
| + s(Longitude) |  |  | <0.001 | 10.3 | 8750.444 |
| + s(Latitude) |  |  | <0.001 | 13.3 | 8728.979 |
| **(b) Age 3** |  |  |  |  |  |
| Intercept | -510.721 | 171.803 | <0.001 |  |  |
| Sex | 12.501 | 4.736 | 0.008 | 0.99 | 8274.399 |
| Latitude | 17.725 | 2.889 | <0.001 | 1.50 | 8272.648 |
| s(Longitude) |  |  | <0.001 | 24.7 | 8083.013 |
| **(c) Age 4** |  |  |  |  |  |
| Intercept | 655.399 | 3.779 | <0.001 |  |  |
| Sex | 20.109 | 5.105 | <0.001 | 1.73 | 11932.84 |
| s(Longitude) |  |  | <0.001 | 6.10 | 11888.33 |
| s(Latitude) |  |  | <0.001 | 8.80 | 11865.64 |

**Figure 2.**
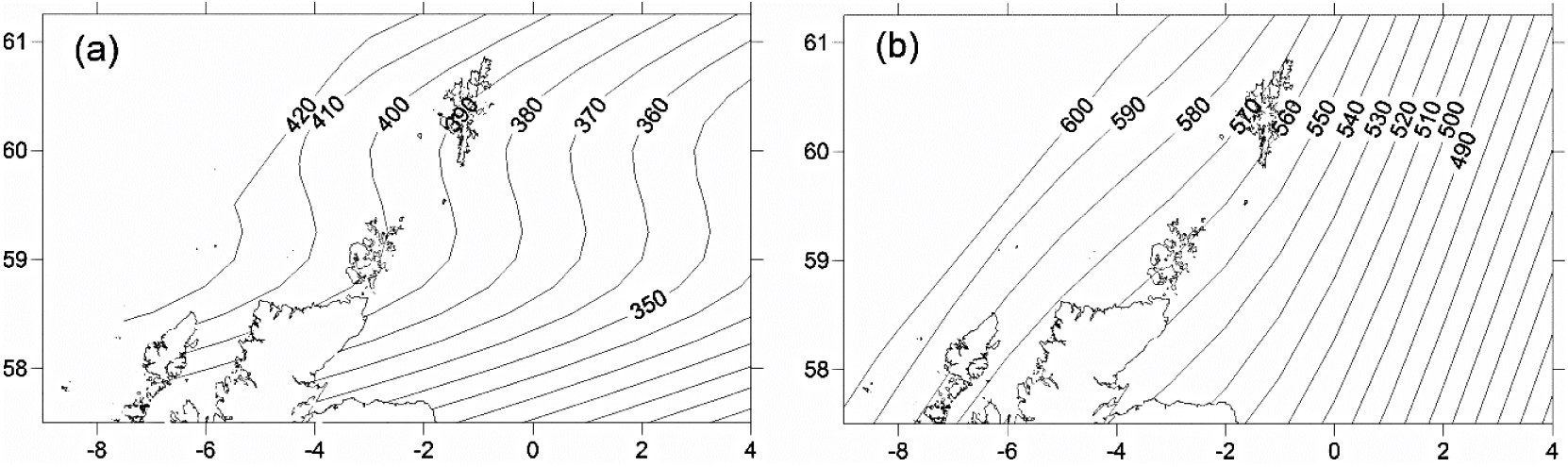
Predicted spatial variation in length at age 2(a) and 3(b) across the study area in 2014.

### 3.2 Spatial variation in length at maturity

Maturity relationships indicated significant effects of length, age, longitude, latitude and period (Table 3). Length at maturity increased from west to east in both females (Figure 3) and males for both study periods. This variation was linked to age, with a predicted 48% of age 2 females mature at 400 mm TL in the far west (ScOW) while none were observed to be mature in the Viking region in the 2013 – 14 period. Similarly, a predicted 80% of age 2 males were mature at 400 mm TL in the far west (ScOW) compared to 23% in the Viking region in the 2013 – 14 period. A slightly higher proportion of males and females were mature at length and age in the later period but there was no significant interaction between period and longitude in females (p = 0.46) or males (p = 0.07) indicating a similar spatial trend over time. A comparison of maturity relationships for autumn and spawning females in 2013 – 14 found no effect of season (p = 0.53) but with similar significant effects of length, age, longitude and latitude to the period model (Table 4). The spatial trend in female maturity for just the genetic samples in 2013 – 14 was consistent with that evident from all surveyed sources, with later and larger size at maturity in the Viking region (Supplement Table S1).

**Table 3.**
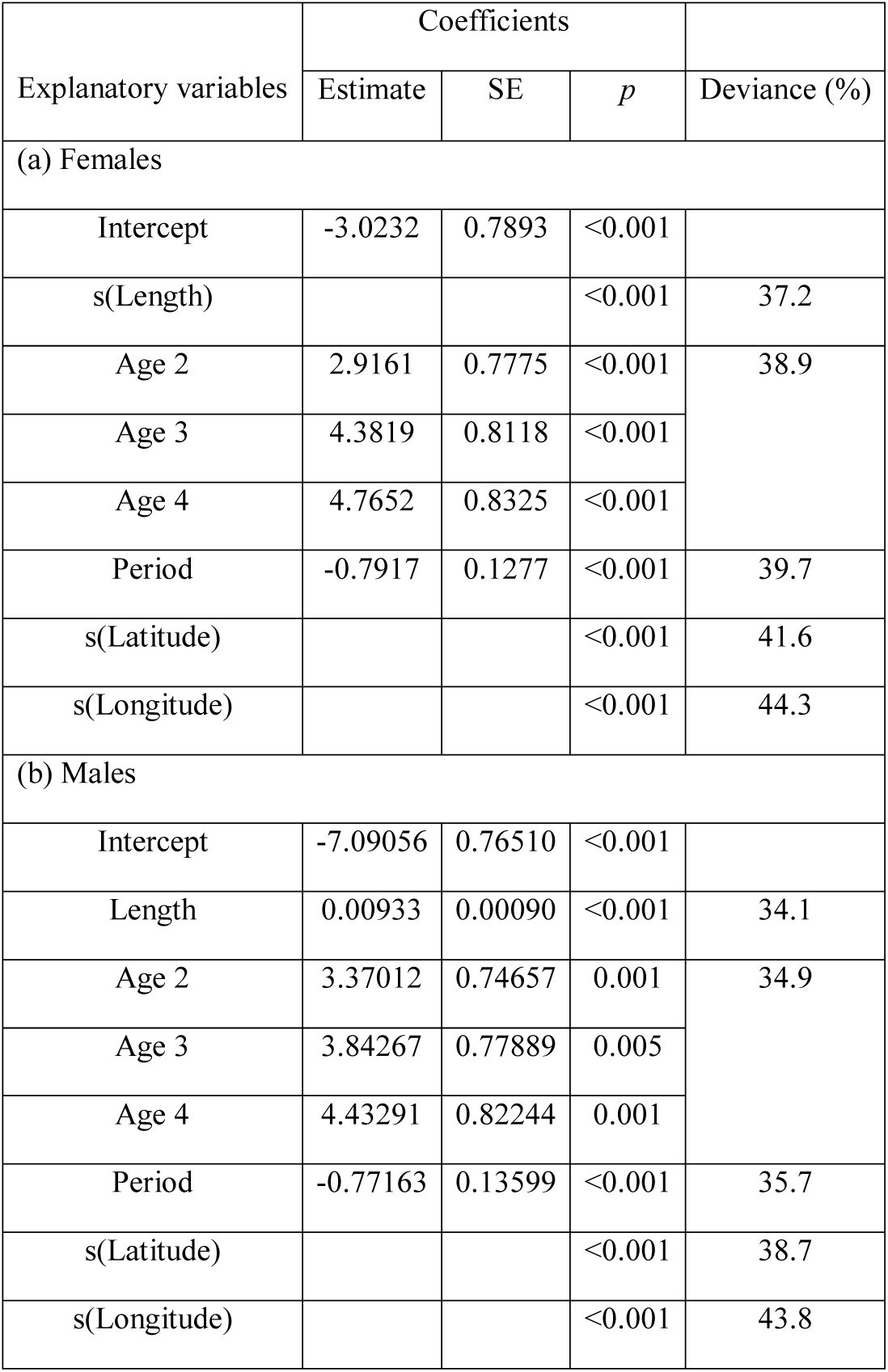
GAM model coefficients and deviance explained for sex-related maturity in relation to length, age, period (decade), longitude and latitude.

**Table 4.** Minimum adequate GAM model coefficients and deviance explained for female maturity in relation to length, age, longitude and latitude for the comparison between seasons.

| Explanatory variables | Coefficients |  |  | Deviance (%) |
| --- | --- | --- | --- | --- |
|  | Estimate | SE | *p* |  |
| Intercept | -6.4245 | 0.9096 | <0.001 |  |
| Length | 0.1156 | 0.01946 | <0.001 | 34.2 |
| Age 3 | 0.8815 | 0.3308 | <0.001 | 35.5 |
| Age 4 | 2.1503 | 0.5959 | <0.001 |  |
| s(Longitude) |  |  | <0.001 | 39.5 |
| s(Latitude) |  |  | <0.001 | 42.1 |

**Figure 3.**
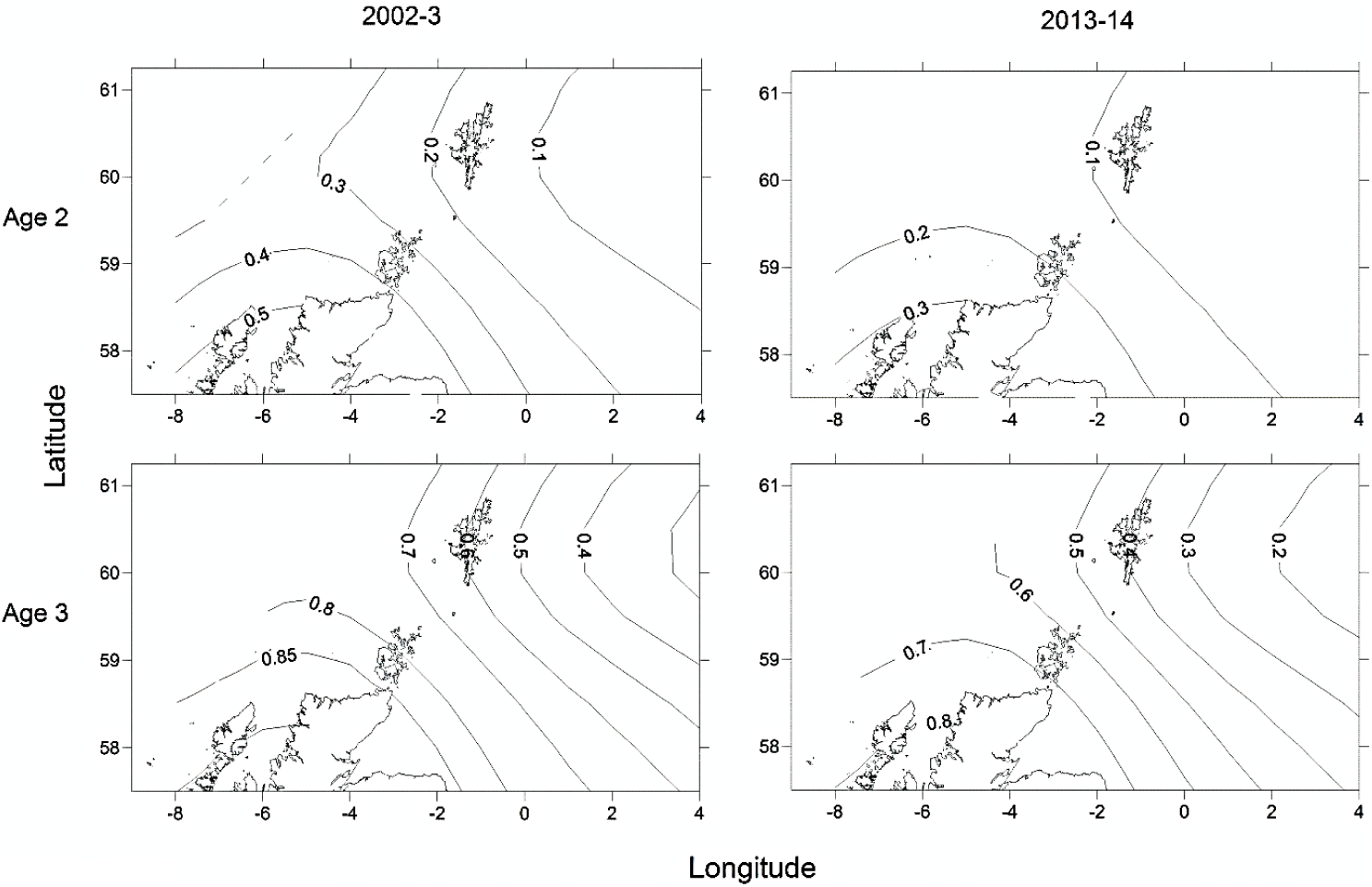
Predicted spatial variation in proportion mature for an age 2, 40cm and an age 3, 50cm female for the two sampling periods.

### 3.3 Evidence of spatial and temporal population structuring

No SNP loci showed consistent deviations from Hardy-Weinberg equilibrium or linkage disequilibrium across populations (Table 1). Outlier analysis identified one loci (SNP_8) as a candidate for directional selection within the dataset, this was retained for the population structure analysis. When the dataset was segregated into the four discrete sampling periods, hierarchical AMOVA analysis revealed significant values at all levels (among periods, among populations within periods and within populations) (Table 5). The differentiation observed among populations within periods was far greater than among periods themselves, implying a temporal stability in the population structuring. Global Fst over all SNPs was low but statistically significant (0.045, *P* = 0.001), with 36% and 33% of all pairwise comparisons within the contemporary and historic datasets respectively being significant. Following sequential Bonferroni correction for multiple comparisons this was reduced to 11% and 17% respectively (Table 6). There was clear and consistent evidence of longitudinal differentiation with ScIW and Viking locations being differentiated (Fst 0.032 – 0.1371) during the spawning and autumn seasons in both the contemporary and earlier datasets. The presence of this longitudinal cline is corroborated by the STRUCTURE analysis for all datasets (Figure 4).

**Table 5.**
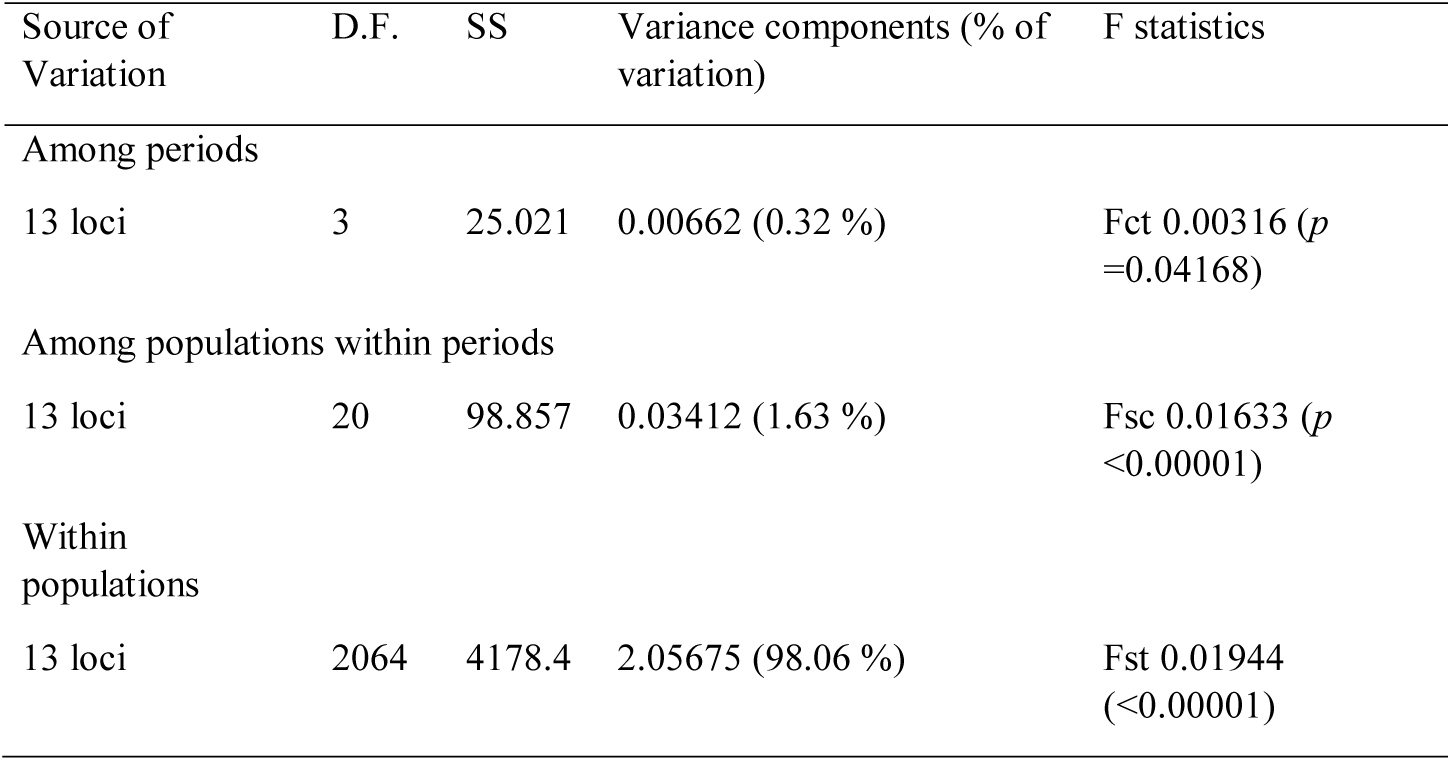
Hierarchical AMOVA based on sampling periods.

**Table 6.**
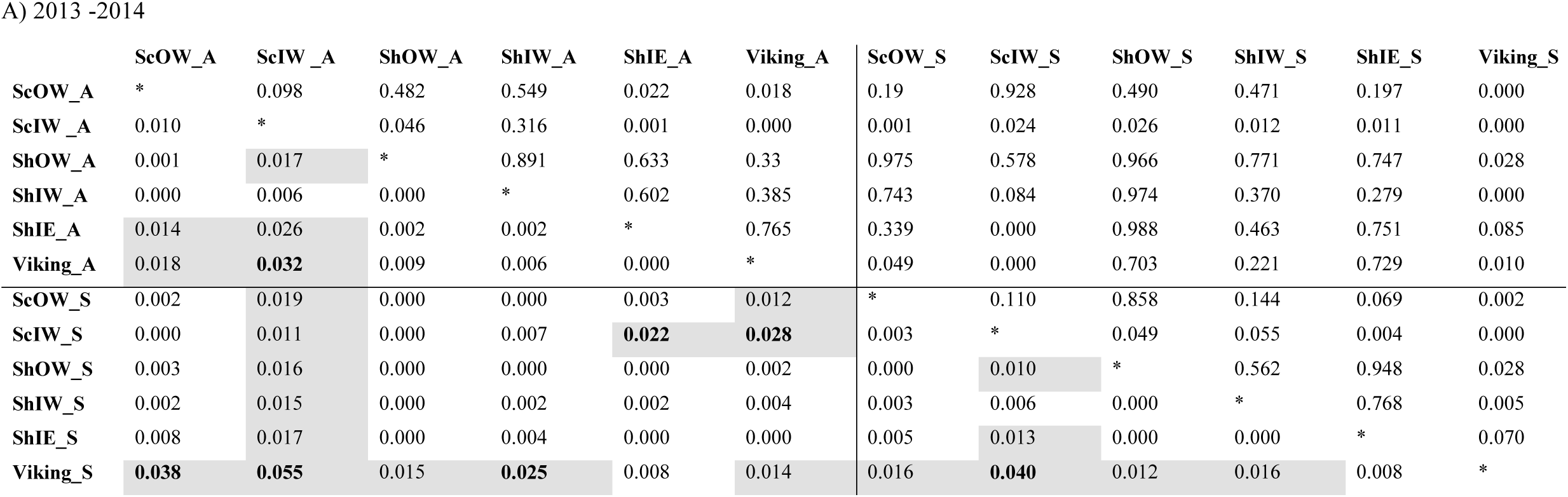
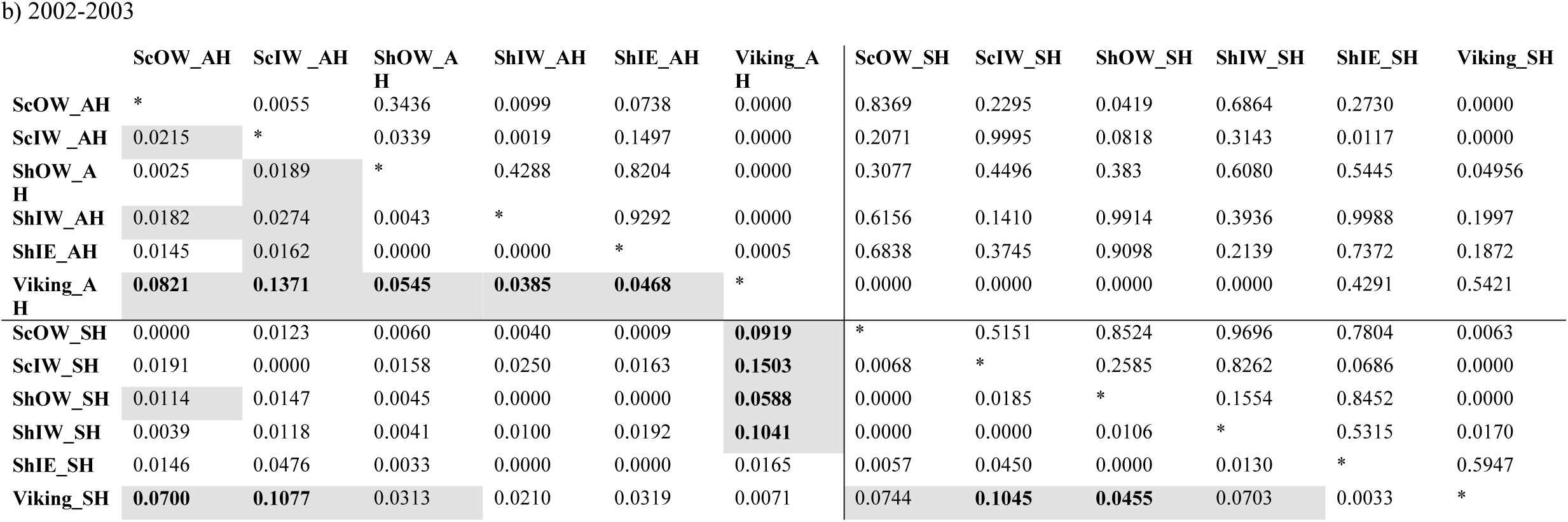
Genetic differentiation between FISA sample locations during a) 2013-2014, b) 2002-2003: pairwise Fst (below diagonal) and *P-value* after 1000 permutations (above diagonal). Pairwise Fst values shaded in grey have a P-value <0.05 and bold indicates significant difference follow sequential Bonferroni correction. (*Location*_A indicates autumn sample while *Location*_S indicates spawning season sample).

|  | ScOW_A | ScIW_A | ShOW_A | ShIW_A | ShIE_A | Viking_A | ScOW_S | ScIW_S | ShOW_S | ShIW_S | ShIE_S | Viking_S |
| --- | --- | --- | --- | --- | --- | --- | --- | --- | --- | --- | --- | --- |
| ScOW_A | * | 0.098 | 0.482 | 0.549 | 0.022 | 0.018 | 0.19 | 0.928 | 0.490 | 0.471 | 0.197 | 0.000 |
| ScIW_A | 0.010 | * | 0.046 | 0.316 | 0.001 | 0.000 | 0.001 | 0.024 | 0.026 | 0.012 | 0.011 | 0.000 |
| ShOW_A | 0.001 | 0.017 | * | 0.891 | 0.633 | 0.33 | 0.975 | 0.578 | 0.966 | 0.771 | 0.747 | 0.028 |
| ShIW_A | 0.000 | 0.006 | 0.000 | * | 0.602 | 0.385 | 0.743 | 0.084 | 0.974 | 0.370 | 0.279 | 0.000 |
| ShIE_A | 0.014 | 0.026 | 0.002 | 0.002 | * | 0.765 | 0.339 | 0.000 | 0.988 | 0.463 | 0.751 | 0.085 |
| Viking_A | 0.018 | **0.032** | 0.009 | 0.006 | 0.000 | * | 0.049 | 0.000 | 0.703 | 0.221 | 0.729 | 0.010 |
| ScOW_S | 0.002 | 0.019 | 0.000 | 0.000 | 0.003 | 0.012 | * | 0.110 | 0.858 | 0.144 | 0.069 | 0.002 |
| ScIW_S | 0.000 | 0.011 | 0.000 | 0.007 | **0.022** | **0.028** | 0.003 | * | 0.049 | 0.055 | 0.004 | 0.000 |
| ShOW_S | 0.003 | 0.016 | 0.000 | 0.000 | 0.000 | 0.002 | 0.000 | 0.010 | * | 0.562 | 0.948 | 0.028 |
| ShIW_S | 0.002 | 0.015 | 0.000 | 0.002 | 0.002 | 0.004 | 0.003 | 0.006 | 0.000 | * | 0.768 | 0.005 |
| ShIE_S | 0.008 | 0.017 | 0.000 | 0.004 | 0.000 | 0.000 | 0.005 | 0.013 | 0.000 | 0.000 | * | 0.070 |
| Viking_S | **0.038** | **0.055** | 0.015 | **0.025** | 0.008 | 0.014 | 0.016 | **0.040** | 0.012 | 0.016 | 0.008 | * |

7 b) 2002-2003
|  | ScOW_AH | ScIW_AH | ShOW_AH | ShIW_AH | ShIE_AH | Viking_AH | ScOW_SH | ScIW_SH | ShOW_SH | ShIW_SH | ShIE_SH | Viking_SH |
| --- | --- | --- | --- | --- | --- | --- | --- | --- | --- | --- | --- | --- |
| ScOW_AH | * | 0.0055 | 0.3436 | 0.0099 | 0.0738 | 0.0000 | 0.8369 | 0.2295 | 0.0419 | 0.6864 | 0.2730 | 0.0000 |
| ScIW_AH | 0.0215 | * | 0.0339 | 0.0019 | 0.1497 | 0.0000 | 0.2071 | 0.9995 | 0.0818 | 0.3143 | 0.0117 | 0.0000 |
| ShOW_AH | 0.0025 | 0.0189 | * | 0.4288 | 0.8204 | 0.0000 | 0.3077 | 0.4496 | 0.383 | 0.6080 | 0.5445 | 0.04956 |
| ShIW_AH | 0.0182 | 0.0274 | 0.0043 | * | 0.9292 | 0.0000 | 0.6156 | 0.1410 | 0.9914 | 0.3936 | 0.9988 | 0.1997 |
| ShIE_AH | 0.0145 | 0.0162 | 0.0000 | 0.0000 | * | 0.0005 | 0.6838 | 0.3745 | 0.9098 | 0.2139 | 0.7372 | 0.1872 |
| Viking_AH | **0.0821** | **0.1371** | **0.0545** | **0.0385** | **0.0468** | * | 0.0000 | 0.0000 | 0.0000 | 0.0000 | 0.4291 | 0.5421 |
| ScOW_SH | 0.0000 | 0.0123 | 0.0060 | 0.0040 | 0.0009 | **0.0919** | * | 0.5151 | 0.8524 | 0.9696 | 0.7804 | 0.0063 |
| ScIW_SH | 0.0191 | 0.0000 | 0.0158 | 0.0250 | 0.0163 | **0.1503** | 0.0068 | * | 0.2585 | 0.8262 | 0.0686 | 0.0000 |
| ShOW_SH | 0.0114 | 0.0147 | 0.0045 | 0.0000 | 0.0000 | **0.0588** | 0.0000 | 0.0185 | * | 0.1554 | 0.8452 | 0.0000 |
| ShIW_SH | 0.0039 | 0.0118 | 0.0041 | 0.0100 | 0.0192 | **0.1041** | 0.0000 | 0.0000 | 0.0106 | * | 0.5315 | 0.0170 |
| ShIE_SH | 0.0146 | 0.0476 | 0.0033 | 0.0000 | 0.0000 | 0.0165 | 0.0057 | 0.0450 | 0.0000 | 0.0130 | * | 0.5947 |
| Viking_SH | **0.0700** | **0.1077** | 0.0313 | 0.0210 | 0.0319 | 0.0071 | 0.0744 | **0.1045** | **0.0455** | 0.0703 | 0.0033 | * |

**Figure 4.**
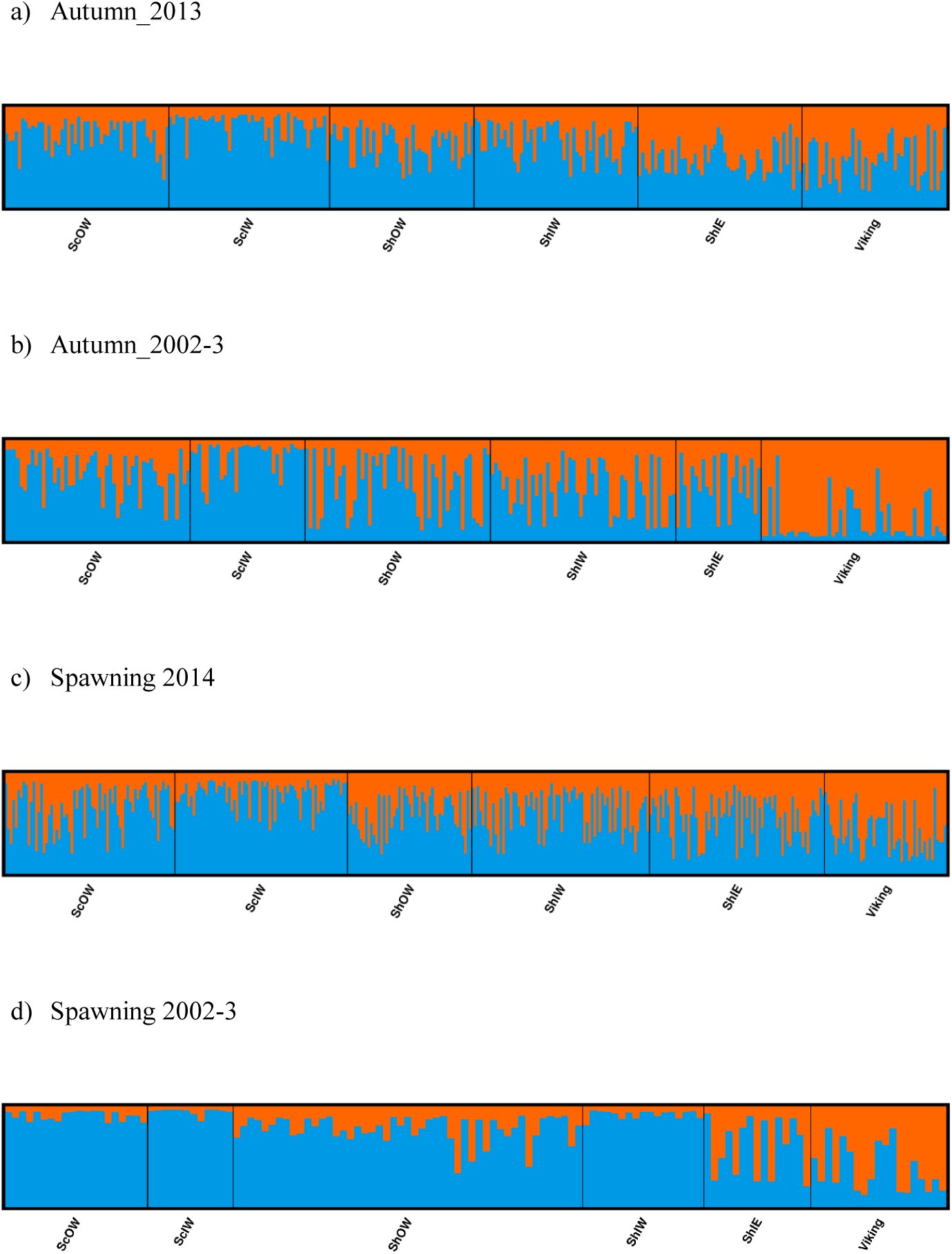
Results from clustering analysis using STRUCTURE with a K value of 2, population groupings are ordered to reflect geographical connectivity with individuals within a given location sorted in ascending longitudinal order for samples captured in (A & B) the Autumn and (C & D) spawning season.

Hierarchical AMOVAs per locus showed that SNP_8, which had been identified by outlier analysis to be a candidate locus under selection, as well as SNP_9 & SNP_10, all showed highly significant differentiation among populations where the average Fst was 0.0974, 0.0441 & 0.0195 per locus respectively. Furthermore, all three loci localised to Linkage group 12 (LG12; Supplementary Table S2). When allele frequency is plotted in relation to sample longitudinal order it is clear that for both SNP_8 and SNP_10 there is a consistent longitudinal cline in allele frequency, which is suggestive of isolation by distance within the study area (Figure 5). A comparison of genetic distance with geographic separation in 2014 was also consistent with an isolation by distance trend (r^2^= 0.30; *P*=0.001; Figure 6), although pairwise Viking and ScIW comparisons were positive outliers with high leverage to this relationship. Conversely, the pairwise comparison between ShIW and ScOW was a negative outlier, and there was no significant difference in pairwise Fst between sites within the region to the west of Shetland except for ScIW (Table 6).

**Figure 5.**
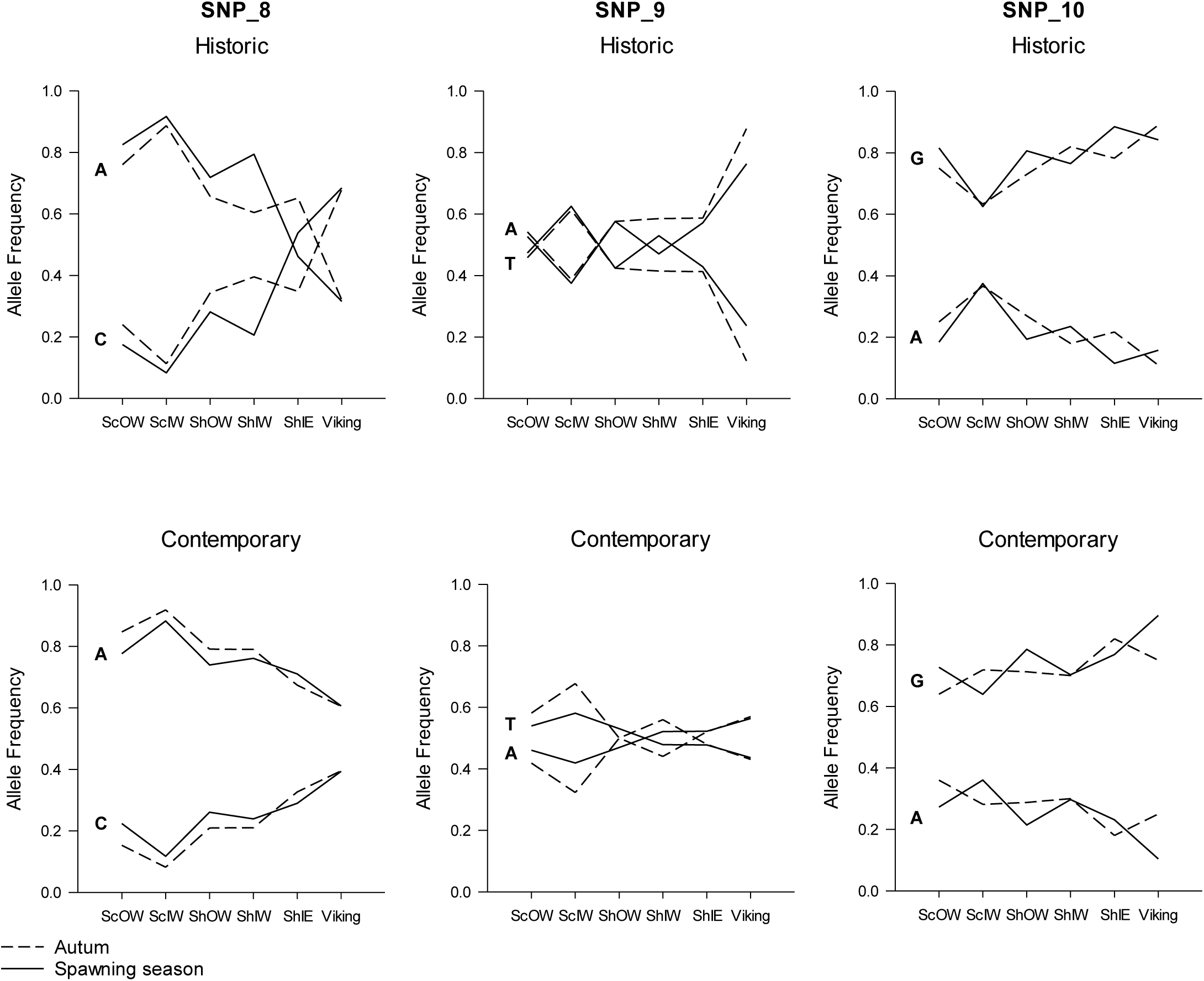
Change in allele frequency in loci SNP_8, SNP_9 and SNP_10 in relation to sample location with the data separated into early (above) and contemporary (below) sample series as well as taken during the spawning season (solid line) and the Autumn (broken line).

**Figure 6.** Isolation by distance plot based on spawning and winter Fst results in Table 6 and geographical separation between samples. Fitted regression line and residual with large leverage (*) are also shown.

## 4. DISCUSSION

The genetic analysis indicates that the North Sea (4) and Scottish west coast (6a) stock units do not correspond to biological populations, and alludes to an even finer scale structuring than that previously reported by Heath et al. (2014). As with recent studies, large F_ST_ differences were related to genomic regions under selection (Berg et al. 2016, Sodeland et al., 2016; Barth et al., 2017), suggesting adaptive genomic divergence while F_ST_ values based on the unlinked neutral SNPs were largely insignificant (data not shown) so truly biological distinct populations were not demonstrated. Studies of adaptive genomic divergence (Barth et al., 2019; Kess et al., 2019) and movements in other Atlantic cod stocks (Robichaud & Rose, 2004; Neuenfeld et al., 2011) suggest two general population types; small coastal resident populations exhibiting year round fidelity and larger populations of seasonal migrants. The temporal persistence of genetic differences within the present study area, coupled to tag based evidence on adult and juvenile movements (Wright et al., 2006 a,b; Neat et al., 2014; Wright et al., 2018), appear to at least partially conform to this pattern.

The three loci which showed highly significant population differentiation, and which included the putative marker under directional selection, were located on Linkage group 12 (LG12). Genome scans of North Sea, western Baltic and fjord-type cod have previously identified high genomic differentiation in inverted regions on LG12 (Sodeland et al., 2016; Barth et al., 2019). This inversion region, in which these three new markers co-locate, has been reported to associate with temperature preference rather than migratory vs non-migratory ecotypes, based on studies from both sides of the Atlantic (Berg et al. 2016, Berg et al. 2017). This association to environmental preference is further rationalised by studies demonstrating the region contains genes associated with temperature and oxygen regulation (Bradbury et al., 2010; Berg et al., 2015). Thus, this inversion could be a prime candidate for a barrier mechanism that might act through either intrinsic (genome incompatibilities) or extrinsic (fitness effect associated with the environment) isolation. The longitudinal trend in allele frequency change found in this study appears to match regional temperature differences, being warm in the west and markedly cooler east of the Shetland Isles in depths > 100 m (see Figure 1). Indeed, electronic tags have found that cod released in the Viking sample area remain in deep, cooler waters all year round experiencing an average temperature of 8.4°C and a range of just 1.5°C, compared to 9.6°C and an annual variation of 7 to 12°C for those released around Shetland and the west coast (Neat et al., 2014).

Genetic differences followed an isolation by distance pattern along a west-east gradient across the study area. Dahle et al. (2018) demonstrated a similar north – south gradient in genetic differentiation of cod along the Norwegian coast, without any distinct change in genetic variation (i.e. allelic diversity or heterozygosity). However, along the Norwegian coast, the barrier to movement is more obvious than in this study, as the cod there inhabit fjords for their entire life-cycle (Knutsen et al., 2018), while in the present study area only cod in Shetland (ShIW) and the coastal waters in the Minch and Lewis (ScIW) show high fidelity as juveniles and adults (Neat et al., 2006; Wright et al., 2006a,b). The strongest evidence for reproductive isolation came from the large positive departures from the isolation by distance trend between Viking and ScIW. Conversely, the lack of significant pairwise Fst differences among samples to the west of Shetland out to the offshore west (ScOW) may suggest a greater exchange, which would be consistent with the wide extent of adult movements in this region (Easey, 1987; Wright et al., 2006a).

There was some evidence of genomic divergence between the west coast cod in coastal waters of the Outer Hebrides and North Rona (ScIW) and those further offshore (ScOW). While further genetic evidence is needed to confirm reproductive isolation, this would be consistent with the limited year round movements evident from tag-recapture studies in the area (Wright et al., 2006a). Movement of young of the year to spawning grounds estimated from otolith chemistry also suggests there is no connectivity between this area and that ∼ 100 kms south (Wright et al., 2006b). Nursery and spawning grounds are found in close proximity to one another in ScIW (Gibb et al., 2007; Wright et al., 2006a) consistent with resident cod populations found elsewhere (Bradbury et al 2011; Grabowski et al., 2011). The temporal stability of this pattern of structuring between samples collected a decade apart further supports the evidence for limited mixing.

Genetic evidence from this study supports previous evidence of a reproductively isolated cod population centred around Viking Bank (Heath et al., 2014; ‘Viking’ deme). Importantly, the more extensive sampling has helped to define the distributional extent of this population, occurring in depths > 100 m up to the northern edge of the continental shelf. However, there was no indication that the Viking group extended west of 1°W during the spawning period. Indeed, the Shetland coastal cod (ShIE) appear to extend onto the western edge of waters > 100 m depth to around 1°W during the spawning period. Given their geographic proximity, it is not surprising that cod tagged with either conventional or archival tags near the east coast of Shetland and near Viking Bank have been found to have overlapping ranges east of Shetland outside the spawning season, although adult Viking cod appear to remain in waters > 100m (Wright et al., 2006b; Neat et al. 2014). Many larval and early juveniles from the Viking area appear to get advected into the Skagerrak from which they make a return migration by the time they mature (Barth et al., 2017; Wright et al., 2018). Based on the disappearance of the majority of age 2 cod from the Skagerrak (Svedäng & Svenson, 2006) and the appearance of high age 2 densities in Viking in the 1^st^ and 3^rd^ quarter IBTS (ICES, 2019a), this return migration is likely to occur around a year before maturation.

The seasonally persistent longitudinal differentiation in both genetic and maturity at size differences provides the first evidence for year round fidelity of the population demes to their spawning areas. This is important as many fish populations, including other cod populations (Neuenfeldt et al., 2013), are known to mix outside the spawning period, which makes it difficult to consider alternative stock boundaries for monitoring and assessment. Year round fidelity can allow for a change in the stock boundaries into management areas that more closely reflect the population structure, as was the case for sandeels (*Ammodytes marinus*) in the North Sea (Wright et al., 2019). The indication of limited seasonal movement of cod in this study is consistent with predicted movement of individual cod based on electronic tagging records (Wright et al., 2006a; Neat et al., 2014). It suggests that the population boundary around 1°W is more appropriate to the scale of population processes than the 4°W line currently used in stock assessment.

The apparent reduction in genetic differences and allele frequency in the contemporary samples compared with that from the previous decade may be evidence of a greater admixture of western and Viking cod in recent times. This suggests some flexibility in population boundaries over time. The late 1990s and early 2000s, when the Heath et al. (2014) samples were obtained, reflected the lowest recorded stock biomass for both North Sea and west of Scotland cod, but there has since been some recovery in the northern North Sea (ICES, 2019,a,b). Past analyses of research vessel survey data have suggested that cod may show density dependent habitat selection (Blanchard et al. 2005) and so it is possible that populations have expanded and with greater overlap between the western and Viking populations of cod.

Cod were larger at age in western waters compared to the deeper northern North Sea. Spatial differences in length at age across the study area appear to persist, based on the lack of a year effect for ages 3 and 4 and the similarity with the findings of Yoneda & Wright (2004). Waters in the west of the study area are the warmest, while coastal waters experience strong seasonal shifts (Berx & Hughes, 2009). Hence, the apparent faster growth in western waters probably reflects a positive effect of temperature, as cod growth in length remains positive up to a peak of 10°C (Righton et al., 2010) and electronic tags have recorded that fish from these waters experience temperatures close to this peak in spring and summer months, with an average annual temperature 1.2°C higher than that experienced by those inhabiting Viking Bank (Neat et al., 2014).

Spatial differences in maturity at length found in this study were consistent with differences between sites to the west of Scotland and Viking Bank detected around the turn of the century (Yoneda & Wright, 2004), but adds a finer spatial resolution that allows for a direct comparison with the scale of genetic variation. The similar geographical pattern of maturity at size and genetic differences indicates there were population level differences in maturation schedules. As temperature just prior to maturation commitment can have a positive influence on the proportion of gadoids maturing (Yoneda & Wright, 2005; Tobin & Wright, 2011), and the deep Viking region is 2°C cooler in the months preceding the autumn equinox (Neat et al., 2014), the observed trend towards later maturation at size moving from west to east would be expected. However, historically the range in maturity-length variation across the North Sea and north west coast of Scotland was much less than that seen since the turn of the century (Yoneda & Wright, 2004). Wright et al. (2011) demonstrated that there was a marked downward shift in the length at maturation in the north west North Sea since the 1980s that could not be explained by warming or reduced competition for resources, whereas no significant change had occurred north east of the meridian. The large difference in maturity at length between Viking and other populations is reflected in maturity at age, with no females maturing at age 2 as was historically the case (Wright et al., 2011). Given the persistence of population differences in maturity-size relationships, it should be possible to use late age at maturity as a phenotypic indicator of whether a cod is from the Viking deme.

The spatial variation in maturity at size at variation can explain published differences between the north east (Wright et al., 2011) and northern North Sea (Heath et al., 2014). Due to uncertainty about the spatial scale of maturity at size variation first reported by Yoneda & Wright (2004), Wright et al. (2011) grouped cod east of the meridian in waters > 100m depth into a north east group in order to examine temporal shifts in the probabilistic maturation reaction norm. This area would have included most of the Viking group evident from this study, and explains the similar length at 50% maturity between that and the present study. In contrast, the length at 50% maturity reported by Heath et al. (2014) in the early 2000s was over 20 cm smaller than that found in this study, which probably reflects the use of the ICES standard roundfish area 1, that extends from 4°W to 4°E (ICES, 2012), which would combine the early maturing coastal cod from Shetland with those from Viking.

Although there was an increase in cod in the northern North Sea from 2006 - 2017, neither stock has fully recovered (ICES, 2019a,b). The lack of any distinct change in the current stock boundaries at 4°W has important implications to current fisheries management. Since 2011, there has been some recovery of cod biomass within the study area including the northern most part of the west coast. In contrast, there has been very little recovery in southern and central parts of the west of Scotland cod stock (Holmes et al., 2014), which, based on past genetic and otolith chemistry, appear to be part of one or more other populations (Wright et al., 2006b; Heath et al., 2014). The spawning stock biomass of the west of Scotland cod is very low according to ICES (2019b), which has led to a zero catch for this stock. Hence, the ability to identify recovery within a population is being compromised by inappropriate stock boundaries.

Past failure to ensure sustainable fishing mortality at a population level appears to be reflected in present day maturation schedules of many cod populations. For example, Olsen et al. (2009) proposed that only the pattern of intense harvesting could explain genetically related fine-scale variation in maturation reaction norms of coastal cod along the Norwegian coast. Differences in maturity at size and age off the north and south coast of Iceland also have been linked to differences in historical patterns of fishing mortality as well as growth (Marteinsdottir & Begg, 2002; Grabowski et al. 2011). In the present study area, fishing pressure was historically much higher in coastal areas (Engelhard et al., 2014) during the period when North Sea coastal populations have undergone large changes in maturation schedules (Wright et al., 2011). Despite intense fishing pressure, the present study suggests there is still a rich population structure, although there is some suggestion from past changes in egg density and areas where spawning adults are no longer recorded that some historic spawning grounds have been lost in Scottish coastal waters (Wright & Rowe, 2019). As small, resident populations tend to inhabit coastal grounds, it is possible that some small components of the stocks have been lost. Other examples of a loss of cod spawning grounds have been reported, such as some in the Gulf of Maine (Ames, 2004). In such coastal areas the spatial scale of genetic differentiation can be very small, as studies of Norwegian coastal cod have demonstrated (Dahle et al. 2018). Hence, the present study highlights the need for measures to conserve population diversity as well as the current focus on stock biomass.

## Acknowledgements

This work was funded by Fishing Industry Science Alliance project 01/13 and Scottish Government project SU01N0, A.D. was funded by a Marine Alliance Science and Technology Scotland PhD studentship. We wish to thank Kenny Coull and Chevonne Angus and all the fish samplers from the Scottish Fishermen’s Federation, Marine Scotland Science and North Atlantic Fisheries College.

## Data Accessibility Statement

Individual sample metadata (location, date of capture, sex, age, genotype) will be made available through the Dryad Digital Repository. SNP sequence information is provided within the supplementary data files.

## Author Contributions

PW jointly designed the study, analysed the phenotypic data and led the writing, A Doyle organised sample collection, performed much of the DNA extraction, genetic analysis and contributed to the writing, JBT performed the RAD Library Construction and contributed to the genetic analysis and writing, AD jointly designed the study, analysed the genetic data and co-led the writing.

