## Supplementary Table 1 for "Linking scales of life-history variation with population structure in Atlantic cod"

**Supplementary table S1:** Loci used for population structure analysis.

| **Locus Name** | **Loci Sequence [SNP annotated]** |
| --- | --- |
| SNP_1 | CATGCTTGTTAAGCAGGATTCCCAAAGCG**[T/C]**ACACAGAGTTCAATGGGGCATGCACGTGGACTGTTTTAAAGTCATCGCAGGCCTTTTAGCTTGATTTCCCGCTGCTTGCACTGGTTCACTTATTTATTTACGTTTTTCTGCCTT |
| SNP_2 | TGCAGGCCAACCTCATCGAGAGCGTCAGCCCAAACACCTTCTGGGAGTGTGTCAACATCGAGAACATAGACCTCTC**[C/A]**ATGAATAGGTACTGGCATGGCAGCATGTGTGAAGCATGTGTGTGATGAGAAAACATAGAATCCTGAG |
| SNP_3 | CATGCAGGCCTACATATAGAAAATACAGAAAACAGA**[T/C]**GGCTCACAGAGAAAAGGTGAACAACCGATTGAAATGACATGGAATCTGAGCGTATTCATAAAATGTCAAACAGTGTGATAAAGTTAAAACCCACGGGTAAGTTATCT |
| SNP_4 | CATGCATTTTTTTAACTCTCT**[A/C]**ATCAAAACAAATAAAAATATCATCCCTCATTTTGACAGCACGCGGCGGTGCTGGAGGATTGGGTCCTTGATGGCTCTATAAGGCTACCAGTCTGCCTTGGTTTGTTTGGACTGCGCTTTGGG |
| SNP_5 | CATGCAAG**[G/A]**CCCCTTGAGAGTTTGACAATTTGTACTTTCTACCTAGCTTTGTGCTATTGTTGCTATGGTCCACTCTGGTCTCCTCATATCACAATGAGTGGATGTTCAGGTGCAGGTCCGACCAGAGCAGAACACGAGCTATAC |
| SNP_6 | CATGCTAATTTCATATATTTGTGCACTTTAATTAGAGCATATTGAGCATATATTTCCATTAGCATGTAGCTTGGTGTC**[G/T]**TACCTTCATGAGGTCGCGGTGGTGTTCCAGGACACTGCAGGTGGTCTGCCAGGTCTGCTGGTAGC |
| SNP_7 | CATGCTTGTATTAAAGAATAAAAAAGAAAACATGAATCCTTGTCGTAATTTTTTCACGCATAATTTTAATAAACAAGATAGTAAG**[A/C]**CCTTGTAAGTCCACATCTGTATTTCATATAGGCCTGTATTAAATACATATAGTTATGA |
| SNP_8 | TGCAGGACTATGAAAGAGCCATCGATTACCATCTCAAACACCTC**[A/C]**TCATCGCCCAGGACCTTGAAGACCGGTACTTACCTCAAACGCAGTGTTTTGTTGTTGTGGTTTCCTCCCAAAGGAGGCTTAGTAACATAGGGAGAATAT |
| SNP_9 | TGCAGGCGAGATGAATCCAAAGAAAAATAAATATACATCATAGTAGTTATCA**[A/T]**GCAACATATATCAATATAGCCATAGAACAATTGTATTAACCCAGTTACTGAGGAGGTTCACGATGAACTTCAAATTCCCATCTTCTTGACC |
| SNP_10 | TGCAGGTGCTCCAGAGCAGGCGCTCCAGAGCTGGCAAACCAGATATTTGCCAGCCTCTTGAGGGAGCCTATACCTTTTGCGAGGCAAAGGGCCCCTCCTGG**[A/G]**CATGCAGGCAGTCAATTCCATCTACCAGGCCCTTGTGGCTTC |
| SNP_11 | CATGCATTCTTTTCTTTAAATGGAGGTATTCAGATGATGAATTATTTCAACGCCAAGTATAGTTTTT**[A/C]**GCCTTTATCTCTAACTTGTGAATGATACCCTTCTTAAGTGTGTTGAAACTTTGTTAAAACTGAGAAACATGTGTGT |
| SNP_12 | TGCAGGGTTTGTGTTAGTGTTGGGGTGGTTATTCAAAAATC**[G/A]**GACCTTGAATTTGGGTATTGTGGTAGCAGCCATCAGAGCATCCTTTCTATCCAAGATGGAAGCGAATCTGTGTTTGATCGCCTGTCATATGAAATAGTAGGC |
| SNP_13 | CATGCCAATGTATCACGTCCACTTATTAAAATACCTGCGACAGC**[G/A]**AAAGTGCTACACAGTGCAAGCTACAAAAAAAATCATCTAAACTATGTTTCACATGCCACACAAAAATATCTAAAGGGGTGTTACCCCATCAAGCGGTCG |

**Supplementary table S2: Minimum adequate GAM model coefficients and deviance explained for female maturity in relation to length, longitude and latitude based on genetic samples only.**

|  | Coefficients | | |  |
| --- | --- | --- | --- | --- |
| Explanatory variables | Estimate | SE | *p* | Deviance (%) |
| Intercept | -5.186 | 1.208 | <0.001 |  |
| Length | 0.136 | 0.022 | <0.001 | 22.9 |
| s(Longitude) |  |  | <0.001 | 43.2 |
| s(Latitude) |  |  | 0.011 | 48.3 |

**Supplementary table S3:** Summary information for the markers used in the study for the complete dataset listing for each loci the global observed heterozygosity (Ho) and unbiased expected heterozygosity (uHe), outputs of hierarchical AMOVA per locus as well as Chromonsome (bold) and positional reference (bp) to which they align to the GadMor3 genome construction.

| **Loci** | **Ho** | **uHe** | **F_ct_** | ***P* - value** | **F_sc_** | ***P* - value** | **F_st_** | ***P* - value** | **Chromosome:** Location |
| --- | --- | --- | --- | --- | --- | --- | --- | --- | --- |
| **SNP_1** | 0.203 | 0.211 | 0.0003 | 0.3765 | -0.0015 | 0.6397 | -0.0012 | 0.6437 | **3** : 10282576 – 10282433 |
| **SNP_2** | 0.194 | 0.196 | -0.0011 | 0.6154 | 0.0044 | 0.1355 | 0.0033 | 0.1759 | **3** : 19406219 – 19406361 |
| **SNP_3** | 0.499 | 0.473 | 0.0100 | 0.0108 | 0.0028 | 0.1777 | 0.0127 | 0.0025 | **4** : 36597569 – 36597426 |
| **SNP_4** | 0.173 | 0.170 | -0.0012 | 0.7992 | -0.0019 | 0.6555 | -0.0031 | 0.7458 | **5** : 6372661 – 6372518 |
| **SNP_5** | 0.212 | 0.234 | 0.0028 | 0.1877 | 0.0063 | 0.1257 | 0.0091 | 0.0520 | **5** : 10640377 – 10640520 |
| **SNP_6** | 0.332 | 0.330 | -0.0007 | 0.6106 | 0.0026 | 0.2117 | 0.0020 | 0.2277 | **8** : 16574032 – 16573889 |
| **SNP_7** | 0.467 | 0.452 | -0.0013 | 0.6216 | 0.0062 | 0.0561 | 0.0049 | 0.0673 | **9** : 24896503 – 24896646 |
| **SNP_8** | 0.362 | 0.367 | 0.0055 | 0.2511 | 0.0925 | <0.00001 | 0.0974 | <0.00001 | **12** : 1174999 – 1175142 |
| **SNP_9** | 0.481 | 0.479 | 0.0112 | 0.0738 | 0.0332 | <0.00001 | 0.0441 | <0.00001 | **12** : 6781635 – 6781492 |
| **SNP_10** | 0.371 | 0.359 | -0.0007 | 0.4677 | 0.0202 | <0.00001 | 0.0195 | <0.00001 | **12** : 17106073 – 17105930 |
| **SNP_11** | 0.428 | 0.431 | 0.0031 | 0.0586 | 0.0001 | 0.4484 | 0.0033 | 0.1864 | **18** : 17368128 – 17368271 |
| **SNP_12** | 0.243 | 0.249 | 0.0008 | 0.3099 | 0.0017 | 0.3522 | 0.0025 | 0.2500 | **21** : 21609862 – 21610005 |
| **SNP_13** | 0.184 | 0.190 | -0.0011 | 0.7885 | -0.0013 | 0.6605 | -0.0023 | 0.7245 | **22** : 17466965 – 17467108 |
